# Metformin and guanylurea reduce survival, but have limited sublethal effects in larval zebrafish (*Danio rerio*)

**DOI:** 10.1101/2023.12.01.569578

**Authors:** S. Williams, W.A. Thompson, N. Masood, O. Birceanu, M. Easwaramoorthy, J. Qiu, J.Y. Wilson

## Abstract

Metformin is the most common first-line oral therapeutic agent used in the treatment of type-2 diabetes, one of the most prevalent chronic diseases in North America. Post excretion, the compound enters wastewater treatment plants where it is partially bio-transformed by bacteria into guanylurea. Both metformin and guanylurea enter freshwater environments in wastewater effluent where they are available for uptake by aquatic biota. However, our understanding of the effects of metformin and guanylurea on aquatic life is limited. We tested the hypothesis that metformin and guanylurea can influence the development of zebrafish (*Danio rerio*), by assessing morphometrics, cardiac development, energetic state, and behaviour of early larvae. Embryos were exposed to environmentally relevant (0.4, 4, 40 μg·L^−1^) and supra-environmental (400 and 4000 μg·L^−1^) concentrations of metformin and guanylurea from the 4-cell stage (3 hours post fertilization; hpf), until first feed (120 hpf). Exposures to 40 μg·L^−1^and higher of both metformin and guanylurea increased mortality. Metformin delayed hatching at the highest concentration tested (4000 μg·L^−1^). The incidence of spinal curvature increased with exposure to both chemicals at supra-environmental levels (400 and 4000 μg·L^−1^ for metformin; 400 μg·L^−1^ for guanylurea). Metformin and guanylurea exposure imposed slight bradycardia in early development, but did not alter oxygen consumption, ATP levels, carbohydrate levels, general swimming, light-dark movement, startle response, or thigmotaxis, irrespective of exposure concentration. The results suggest similar and low sensitivities of larval fish to both metformin and guanylurea. Apart from a small increase in mortality, these compounds impart a modest impact to the early-life stages of zebrafish that are largely limited to supra-environmental concentrations.

## INTRODUCTION

Metformin is a potent antihyperglycemic insulin-sensitizing agent, primarily prescribed to patients with non-insulin-dependent diabetes mellitus. Metformin has quickly become one of the most widely consumed pharmaceuticals worldwide and was added to the World Health Organization’s list of essential medicines in 2013 (WHO 2021). Metformin and other guanidine derivatives, such as phenformin and buformin, are compounds containing a biguanide core, composed of the coupling of two guanidine units: HNC(NH_2_)_2_. For metformin, one of the two guanidine subunits has two attached methyl groups which confer polar basic properties and high stability. Metformin has a low log K_ow_ of -2.63 and in tandem with its high polarity, is freely soluble in polar solvents such as water (Georgita et al. 2018). Under aerobic wastewater treatment conditions, metformin can become bacterially degraded into the transformation product guanylurea (Trautwein and Kümmerer 2011), through the removal of methyl groups from the terminal nitrogen (Markiewicz et al. 2017). Guanylurea is resistant to common wastewater treatment practices, like UV light irradiation and ozonation, and is the major end-product (Trautwein and Kümmerer 2011) in metformin biodegradation. Similarly to metformin, guanylurea is expected to be highly hydrophilic (estimated K_ow_ of -1.22 to -2.22; Straub et al. 2019) and mobile in aquatic environments (Scheurer et al. 2012). Metformin is among the most abundant pharmaceuticals detected in waters receiving wastewater effluent, with surface water concentrations ranging between 0.1-30 µg·L^−1^ (Oosterhuis et al. 2013; Niemuth and Klaper 2015; Littlejohn et al. 2023). Guanylurea is also detected in surface waters, albeit in some cases, at higher concentrations than its parent compound (Scheurer et al. 2012). Despite this, limited work has been carried out describing the impacts of metformin, and its biotransformation product guanylurea, on aquatic life such as fish.

Literature concerning the impact of metformin on fish has produced mixed results. Embryonic zebrafish (*Danio rerio*) exposed to metformin for 120 hours post-fertilization (hpf; 0.129 µg·L^−1^ – 1,291.6 µg·L^−1^) had differences in gene expression (19 upregulated and 3 downregulated) across exposure concentrations, particularly in genes involved in regulating neural and cardiac development (Phillips et al. 2021). Zebrafish exposed to ≥20 µg·L^−1^ of metformin for 120 hpf showed increased incidence of scoliosis and abnormal pigmentation (Elizalde-Velázquez et al., 2021). Velazquez and colleagues (2021) documented increased mortality, malformation rate, hatch rate and oxidative stress in 96 hpf zebrafish exposed to < 100 μg·L^−1^ of metformin. Exposures to metformin (1–100 μg·L^−1^) in Japanese medaka (*Oryzias latipes*) that covered the embryonic to larval stage, found disruptions in the levels of L-lysine, L-proline, and DL-3-aminoisobutyric acid, while increasing steric, palmitic, and arachidonic acid over a 28-day period (Ussery et al. 2018). Metformin exposed fish also had reduced mass at ≥ 3.2 μg·L^−1^ and displayed diminished lengths at 10 μg·L^−1^ L (Ussery et al., 2018) suggesting that metformin may disrupt growth and development. Indeed, studies have demonstrated that metformin exposure from 4 hpf to 9 months of age can possibly impair ATP production by reducing COX1 activity (Barros et al. 2022). Conversely, life cycle exposures of fathead minnow to metformin have shown no impairment on growth metrics (Parrott et al. 2021; Parrott et al. 2022). Behaviorally, metformin exposure can lead to perturbations in function (Philips et al., 2021); other studies report no clear impact on locomotion following exposure (Godoy et al. 2018; Jacob et al. 2018)

Guanylurea exposure in fish has received comparatively less attention than that of metformin. *Oryzias* larvae exposed to increasing concentrations of guanylurea (1.0-100 μg·L^−1^) for 28 days had decreased length and weight (Ussery et al. 2019). A related study increased the duration of the exposure from 28 days to 165 days post-fertilization (dpf) but there was no effect on adult (165 dpf) medaka body size (length, wet weight, or condition factor; Ussery et al., 2019). In two experiments conducted by Jacob and colleagues (2019), embryos and nine-month-old juvenile brown trout (*Salmo trutta*) were exposed to increasing concentrations of guanylurea (0, 100 and 1,000 μg·L^−1^) for three weeks. Guanylurea exposure did not alter endpoints in larvae (heart rate and hatch time) or juveniles (body weight, length, stress proteins and lipid peroxidases; Jacob et al., 2019). The conflicting results with regards to larval health after metformin and guanylurea exposure may be the result of differences between species responses, but clearly further work is required to identify sublethal impacts of exposure to aquatic life.

This study tested the hypothesis that metformin and guanylurea would pose a risk to the early development of larval zebrafish. Early-life stages present a susceptible window for contaminants, as alterations in controlled developmental plans may detrimentally impact the resulting function of larvae (Thompson et al., 2017). Static renewal exposures over 5 days were used to assess the effects of metformin and guanylurea on cumulative mortality (120 hpf), hatching (48 hpf), morphology (72 hpf) and percent abnormality (120 hpf), heart rate (24, 48 and 72 hpf), energetics (oxygen consumption, ATP, glucose, and glycogen; 120 hpf), and behavior (thigmotaxis, light/dark responsiveness, and startle response; 96-120 hpf). The results of this study provide important data for understanding the effects of metformin and guanylurea on developing fish and may be useful for risk assessment of pharmaceuticals in aquatic environments.

## MATERIALS AND METHODS

### Test chemicals

Metformin hydrochloride (1,1-dimethylbiguanideine hydrochloride; CAS# 1115-70-4) was purchased from Toronto Research Chemicals (Toronto, Ontario, Canada). Guanylurea (CAS# G844495) was purchased from Sigma-Aldrich (Oakville, Ontario, Canada).

### Fish Care and Breeding

Breeding stocks of mature TL wild-type zebrafish, age 4-12 months, were maintained in a semi-recirculation rack housing systems with 15% daily distilled water replacement and automated sodium bicarbonate and salt dosing (Instant Ocean, SpectrumBrands, USA). The system was maintained at 28.5°C, pH of 7-8, and conductivity of 450 -470 μS with a light-dark cycle of 14:10 hours. The fish were fed two times a day with Nutrafin MAX Tropical Flakes (Tetra, USA) and live artemia (GSL Brine Shrimp, US). All fish care was provided in accordance with McMaster University’s animal ethics research board using approved protocols under Animal Use Protocol #20-06-23.

Embryo traps were inserted into a tank the night prior to embryo collection. Traps with embryos were removed 1.5 hours after first light, carefully rinsed to remove all debris, bleached (0.01% in E3 embryo media [5mM NaCl, 0.17mM KCl, 0.33 mM MgSO_4_, and 0.33 mM Cacl_2;_ pH 7.0-7.5] for 1 minute, rinsed with embryo media, and placed in petri dishes (∼100 individuals per dish) with E3 embryo media. Embryos were observed under light microscopy to ensure only 12-32 cell stage embryos were selected for exposures.

### Analysis of exposure medium via LC-MS

Metformin and guanylurea exposure solutions were quantitatively assessed using LC-MS with a 1290 Infinity HPLC s. Samples of 20 mL exposure were randomly sampled before carrying out a 50% water change out using a filtered sterile syringe (Henk Sass Wolf Normject, 4200-X00V0 20 mL) with a glass microfiber disposable syringe filter (Whatman; pore size 0.7μm, 6890-250725 mm GD/X). Filtered samples were stored in 20-mL scintillation vials. The LC-MS-MS analyses were performed using a WatersXevo TQS mass spectrometer coupled to an Acquity ultra-performance liquid chromatography system.

### Exposures, mortality, and hatching

Fish embryo acute toxicity tests with *D. rerio* were based off of the OECD guideline 236 (OECD, 2013). ∼100 newly fertilized embryos (2 hpf) were transferred to petri dishes (100 x100 mm) filled with 100 ml of metformin or guanylurea dissolved in E3 medium. Metformin and guanylurea solutions were diluted from one of two stock solutions (4 mg·L^−1^ for the 0.4, 4 μg·L^−1^ treatment groups and 40 mg·L^−1^ for the 400, 4000 μg·L^−1^ treatment groups). Control plates containing E3 medium were used for controls. A total of 12 independent exposure experiments were carried out and each experiment had 3 replicate petri dishes for each concentration.

Embryos were kept at 28.5 °C and monitored daily for survival and developmental milestones for the duration of the experiment with periodic removal for endpoint collection. Chorions (post-hatching) and dead embryos were removed daily. Control and test solutions were renewed daily after observation of embryos under stereomicroscopy (Zeiss [Jena, Germany], model Axio Zoom. V16); at least 40% of the volume of each petri dish was replaced with pre-made exposure water. Mortality and hatching were observed every 24 hours. General hardness (GH), carbonate hardness (KH), pH, nitrite (NO_2_^-^), and nitrate (NO_3_) were not different between groups.

### Larval fixation and morphological and abnormality assessment

Zebrafish larvae were euthanized at 72 hpf and 120 hpf with tricaine methane sulfonate (300 mg·L^−1^) buffered with sodium bicarbonate (mg·L^−1^). For fish at 72 hpf, tricaine was replaced with 1 mL neutral buffered formalin for 1 hour for larval fixation. Post fixation, larvae were washed with 2 mL phosphate buffered saline (PBS), dehydrated with 1 ml 70% ethanol, and stored at 4°C for storage. A Zeiss Axio Zoom V16 (Zeiss, Jena, Germany) was used to acquire 60 (3 replicates of 20) lateral 4752-pixel images of the embryos per concentration. Seven morphological parameters were measured in images from 72 hpf larvae using the free graphical image analysis software ImageJ (National Institutes of Health, Bethesda, MD, USA) with the observer blind to treatment group: Standard length (SL), snout-vent length (SVL), height at anterior of anal fin (HAA), head depth (HD), and eye diameter (ED), yolk sac depth and yolk sac length. The calculation of yolk sac volume was done using the following formula according to Subhan et al., 2020: V= π/6 • L• H^2^, where V = Yolk sac volume (mm^3^) L = Yolk sac length (mm), and H = Yolk sac depth (mm). Measurements of morphology were only carried out in animals that did not exhibit abnormalities.

At 120 hpf, larvae (total 60-80 larvae taken from both respirometry trials [described later], and unused from petri dishes) were euthanized and deformities of the spine (scoloiosis, bends, or kinks), yolk sac edema, and pericardial edema were observed under aide of microscope. Percentages were calculated from the total number of each type of malformation observed/total larvae of each replicate (n=12).

### Larval behavior

All behavioral assays were completed using the Danio vision observation chamber (Noldus B.V., Wageningen, the Netherland) and EthoVision XT version 15 video tracking software. Analyses were performed in an isolated room and the chamber was maintained at 28.5 °C. Briefly, larvae were transferred to clear 96-well plates with lids and allowed to acclimate overnight (96 hpf at time of measurement) before analyses for visual motor and acoustic startle responses (300 μL test medium). Larvae were transferred to a 24 well plate and were allowed to acclimate overnight (120 hpf at time of measurement) for general locomotor activity and thigmotaxis (2 ml test medium). In zebrafish, visual motor response is a method used to measure locomotion behavior in alternating periods of light (lower activity) and dark (higher activity; Thompson et al., 2017). This assay is used to measure both locomotor activity and behavioral response to different light conditions, exposing larvae to light or dark periods. The assay ran for 60 minutes over four alternating periods of light (maximum light intensity) and darkness lasting 7.5 min (450 s) each. Data was collected in 3 trials (16 larvae/treatment group/plate) for a total of 48 larvae per treatment. Startle response in zebrafish involves an evoked “fast start” behavior consisting of a rapid turn and swim away from the stimulus (Rice et al. 2011). After a 5-minute acclimation period, larval zebrafish were subjected to 10 acoustic/vibrational stimuli (Danio vision intensity setting 6) with a 20 s inter-stimulus interval. The variable of interest used as a proxy for startle response was *maximum velocity* (mm/s), which was collected as the maximum speed in the first second after the implementation of the stimulus. General locomotor activity and thigmotaxis were collected in 3 trials (16 larvae/treatment group/plate) for a total of 48 larvae per treatment. General locomotor activity was determined over 30 minutes in a reduced light within the Danio vision chamber (20% of total maximum intensity); activity was measured as the total distance moved (mm) over the whole arena (i.e., each well). Thigmotaxis determined the time (seconds) spent in the inner and outer areas of the well giving an estimation of the anxiety-like behaviour of the animal (Thompson and Vijayan 2020a). The outer area was defined as within 4 mm of the wall and the total area for inner and outer areas were equivalent. Thigmotaxis was calculated as the percent time spent in the outer zone.

### Oxygen consumption and heart rate

Oxygen consumption rates (MO_2_) were determined at 28 ± 0.1°C for larval zebrafish at 5 dpf. Prior to assessment, 4 mL vials (15 x 48; D x H), with integrated oxygen sensors were calibrated using air saturated water at each concentration tested (for both metformin and guanylurea). A total of 4 sensors were connected to the Pyro science Firesting-O_2_ (4 channel) system. Larval zebrafish were added into 3 vials (∼40 fish/vial), with one vial remaining empty to account for any bacterial respiration of the system water. Following loading of the first chamber, screw caps with a septum were used to seal the chambers, any visible bubbles were removed, and vials were immediately placed into a water bath maintained at 28.5°C. Upon the first vial entering the water bath, data logging software (Pyro Workbench) was used to collect oxygen measurements in 5 s intervals. Closed oxygen respiration assessments were conducted for 1.5 h. Three replicates of each experimental concentration were collected from 4 separate exposure cycles of zebrafish, equating to a total n of 12 for this assessment. Following completion of the assessment, the total number of larvae were confirmed. Analysis of oxygen consumption was carried out using Pyro Data Inspector software, with MO_2_ calculated as the oxygen consumed between 20-50 min of the assessment. The first 20 minutes were removed to account for the variability noted with intrusion of the water bath. The rate of the blank chamber was subsequently subtracted from all slopes calculated from fish containing chambers. Oxygen consumption was normalized to the total number of larvae used for each assessment (µmol O_2_/larvae/h). Heart rate was determined by direct observation under light microscopy, counting the number of beats observed for 15-30s and multiplying the heart rate collected to equate to 1 min of data collection (eg. 30 s data was multiplied by 2). Data was collected in 3 trials (15 larvae/treatment group/plate) for a total of 45 larvae per treatment. Each trial was temperature controlled at 28 ± 0.7°C.

### ATP, glucose, and glycogen

At 120 hpf, zebrafish were euthanized in MS-222, and groups of 15 larvae from each treatment were flash frozen in liquid nitrogen and stored at -80°C until assessment (n=6). Larval samples were sonicated (Sonifier cell disruptor 350; Branson, USA) in a protease inhibitor cocktail (in 50 mM Tris-Buffer; pH 7.5; Roche diagnostics, USA), and centrifuged for 2 min at 13,000 x g. Supernatant was divided to either measure ATP, free glucose, or to undergo glycogen digestion via amyloglucosidase. Measures of ATP, glucose, and glycogen were carried out as described previously (Thompson et al., 2023).

### Statistical Analysis

All statistical analyses were performed using Prism 8.0 (graphpad). Data sets were analyzed by either a one-way ANOVA, repeated measures two-way ANOVA (in the case of the acoustic tap response with tap number and treatment as main effects), or in the case of violations of normal distribution and homoscedasticity, a Kruskal-Wallis one-way ANOVA and Dunn test was employed. All data are presented as mean ± standard deviation.

## RESULTS

### Cumulative Mortality and hatching success

The cumulative mortality (3-120 hpf) in the control groups for metformin and guanylurea experiments were 10% and 15%, respectively (Figure 1). Exposure to metformin significantly increased cumulative mortality in the 40, 400, and 4000 μg·L^−1^ treatment groups (Kruskal-Wallis; DF = 5, H = 29.35, p < 0.01; Fig. 1A). Guanylurea exposure significantly increased mortality in the 40 and 4000 µg · L^-1^ treatment groups (Kruskal-Wallis; DF = 5, H =17.48, p = 0.0037; Fig. 1B). Exposure to 4000 μg·L^−1^ metformin delayed hatching at 48 hpf (Kruskal-Wallis; DF= 5, H=20.95, p=0.0008; Fig. 1C). Hatching was not altered by exposure to guanylurea (Kruskal-Wallis; DF=5, H=1.725, p>0.05; Fig. 1D).

**Fig 1.**
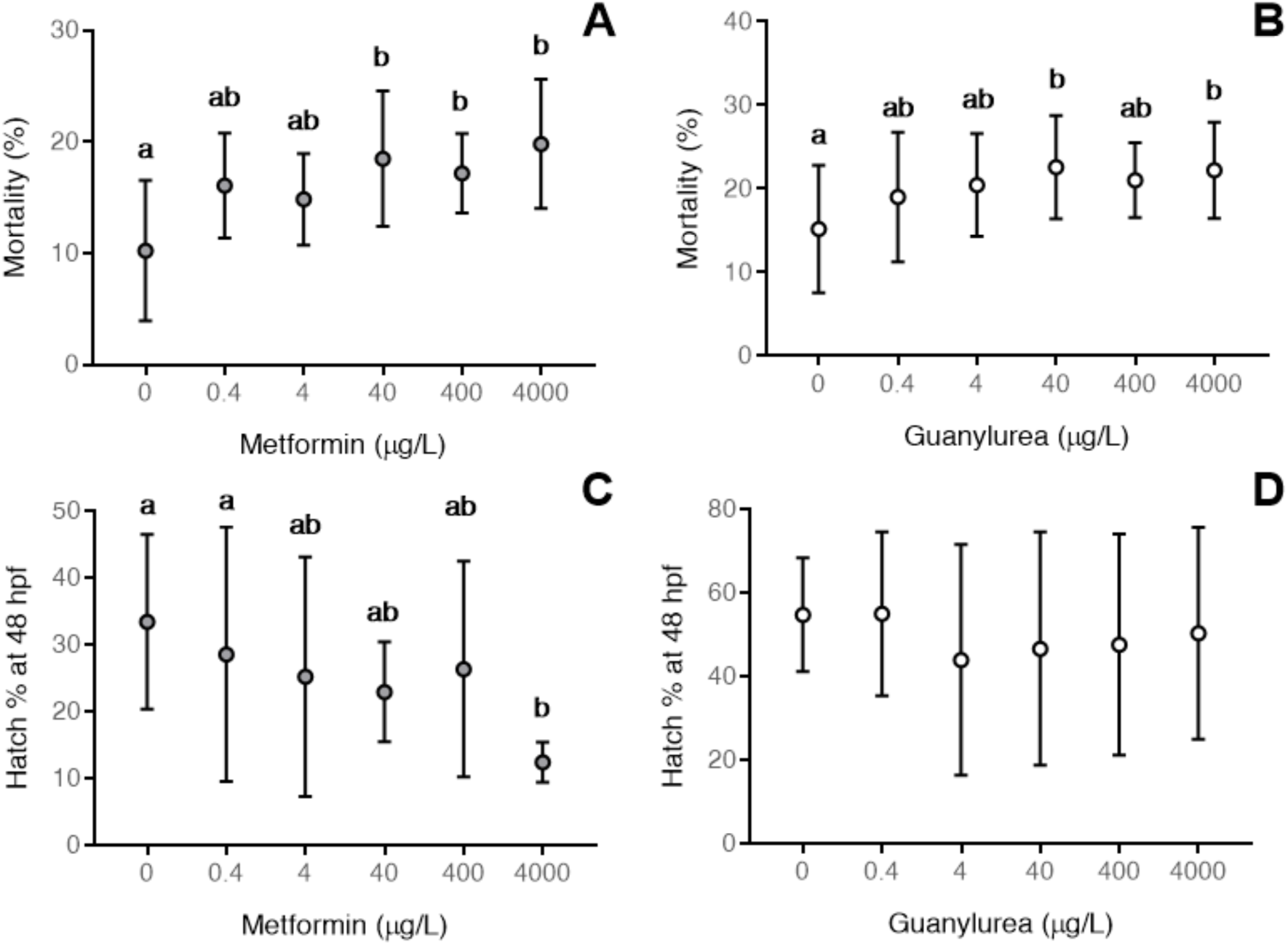
Metformin and guanylurea increase mortality, but only metformin delays hatching. Cumulative mortality in zebrafish exposed to increasing concentrations of (A) metformin and (B) guanylurea. The hatching at 48 hpf of zebrafish exposed to (C) metformin and (D) guanylurea. Values are displayed as mean ± standard deviation. Different letters denote significant differences between groups (n=21 for mortality; n=12 for hatch; each replicate is a dish of ∼100 individuals)

### Abnormalities and Morphometrics

Exposure to 400 and 4000 μg·L^−1^ metformin increased prevalence of spinal abnormalities (ANOVA DF = 5, F =4.704, p = 0.001; Fig. 2A). Metformin exposure did not alter the prevalence of yolk sac (ANOVA DF = 5, F = 1.39, p>0.05; Fig. 2E) or pericardial edema (ANOVA DF=5, F=1.38, p>0.05; Fig 2C). Guanylurea (0.4 compared to 4000 μg·L^−1^ only) increased the incidence of spinal curvature in zebrafish larvae (ANOVA DF=5, F=3.6, p=0.006; Fig. 2B). Cardiac edema did not increase following exposure to guanylurea (ANOVA DF=5, F=2.02, p=0.0874; Fig 2D), nor did the incidence of yolk sac edema increase (ANOVA DF=5, F=1.387, p=0.241; Fig 2F). Metformin exposure decreased snout-vent length (Kruskal-Wallis =36.24, df=5, p-value=<0.01) at the highest tested concentrations (400 and 4000 μg·L^−1^; Table 1). Height at anterior anal fin (Kruskal-Wallis =22.3, DF=5, p-value=<0.01) and eye diameter (Kruskal-Wallis =33.66, DF=5, p-value=<0.01) were decreased by metformin but only at the highest tested concentration tested. Standard length (Kruskal-Wallis =44.1, DF=5, p-value= 0.21), yolk sac area (Kruskal-Wallis =36.43, DF=5, p-value= 0.3) and head depth (Kruskal-Wallis=34.1, DF=5, p-value= 2.2) were not impacted by metformin exposure Exposure to guanylurea decreased standard length in all tested concentrations (Kruskal-Wallis =44.16.97, DF=5, p-value= <0.01). Snout-vent length was decreased in all tested concentrations except for 4 µg · L^-1^ (Kruskal-Wallis =43.97, DF=5, p-value=<0.01). Eye diameter decreased with guanylurea at the highest tested concentration ^(^ANOVA DF = 5, F =2.41, p = <0.04). Yolk sac area (Kruskal-Wallis =34.1, DF=5, p-value= 2.2), head depth (Kruskal-Wallis=2.32, DF=5, p-value= 0.8), and height at anterior anal fin (Kruskal-Wallis =6.89, DF=5, p-value= 0.2) were not altered by guanylurea exposure.

**Fig 2.**
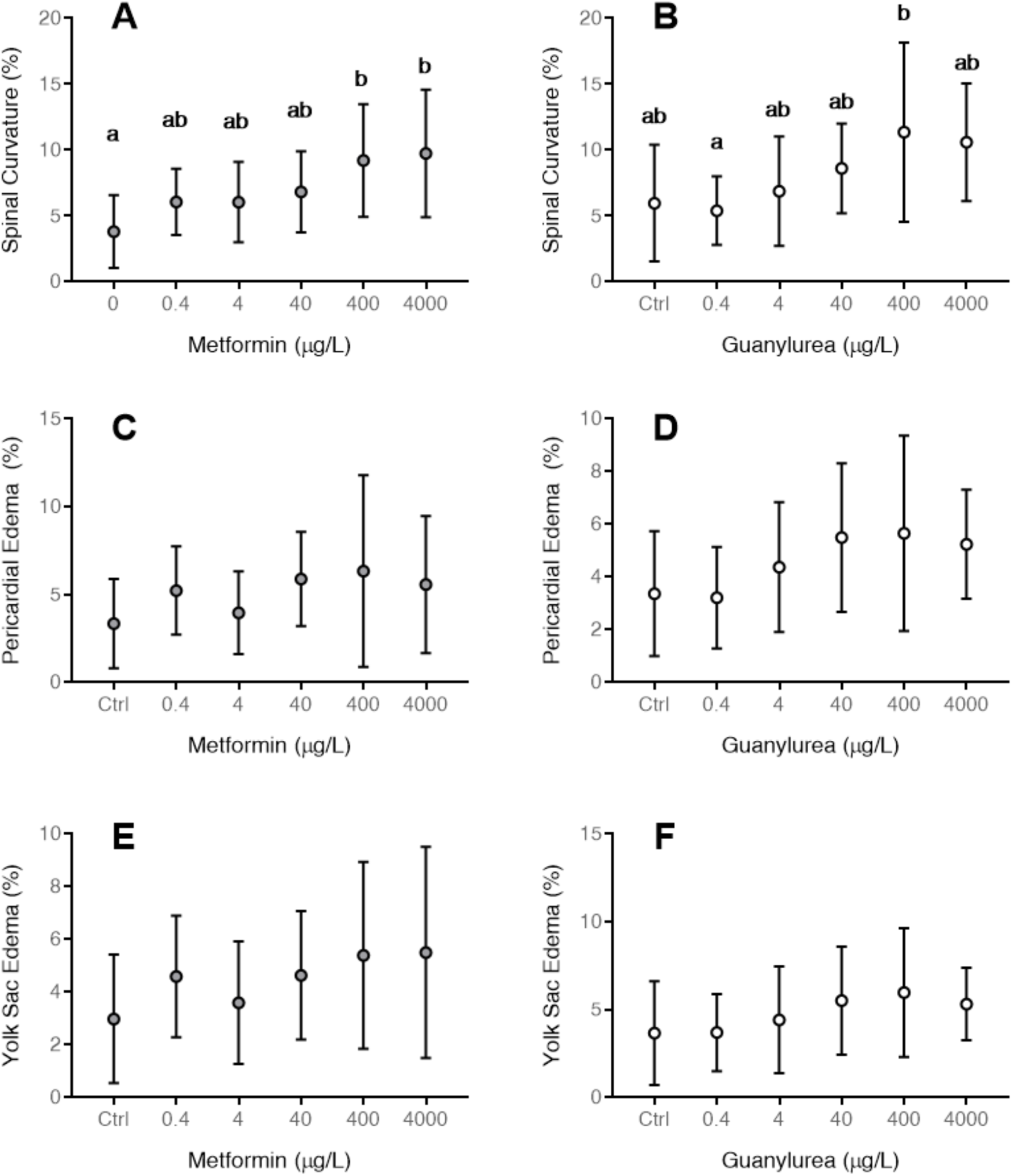
Supra environmental levels of metformin and guanylurea increase the incidence of spinal curvature. Percentage of zebrafish larvae at 120 hpf presenting with spinal curvature, pericardial edema, and yolk sac edema following exposure to metformin (A, C, and E, respectively) and guanylurea (B, D, and F, respectively). Values are displayed as mean ± standard deviation. Different letters denote significant differences between groups (n=12; observations made from 60-80 larvae per replicate).

**Table 1:** Morphometrics of embryo-larval zebrafish exposed to metformin and guanylurea for up to 120 hpf. Data are presented as treatment means ± standard deviation. Morphological structures were determined in lateral images. Values with an alphabetical superscript in common are significantly different (p < 0.05, n=13-16 for standard length; n=60 for other measures). SL= standard length; SVL= snout-vent length; YSA= yolk-sac area; HAA= height at anterior anal fin; HD= head diameter; ED= eye diameter.

| Treatment | SL | SVL | YSA | HAA | HD | ED |
| --- | --- | --- | --- | --- | --- | --- |
| Metformin<br>($\mu\text{g/L}$) | | | | | | |
| 0 | $3.4 \pm 0.01$ | $1.99 \pm 0.08^a$ | $7.5 \pm 0.1$ | $0.27 \pm 0.1^a$ | $0.50 \pm 0.63$ | $0.33 \pm 0.63^a$ |
| 0.4 | $3.5 \pm 0.03$ | $1.93 \pm 0.09^{ab}$ | $7.4 \pm 0.7$ | $0.27 \pm 0.09^a$ | $0.51 \pm 0.45$ | $0.32 \pm 0.45^a$ |
| 4 | $3.4 \pm 0.03$ | $1.94 \pm 0.04^{ab}$ | $7.4 \pm 0.7$ | $0.26 \pm 0.09^a$ | $0.49 \pm 0.41$ | $0.32 \pm 0.41^a$ |
| 40 | $3.4 \pm 0.07$ | $1.95 \pm 0.1^{ab}$ | $7.2 \pm 0.7$ | $0.27 \pm 0.1^a$ | $0.5 \pm 0.2$ | $0.32 \pm 0.17^a$ |
| 400 | $3.4 \pm 0.04$ | **$1.97 \pm 0.1^a$** | $7.3 \pm 0.7$ | $0.27 \pm 0.2^a$ | $0.51 \pm 0.4$ | $0.33 \pm 0.29^a$ |
| 4000 | $3.4 \pm 0.01$ | **$1.90 \pm 0.17^c$** | $7.4 \pm 0.6$ | **$0.25 \pm 0.01^b$** | $0.48 \pm 0.34$ | **$0.30 \pm 0.45^b$** |
| Guanyldurea<br>($\mu\text{g/L}$) | | | | | | |
| 0 | $3.37 \pm 0.24^a$ | $1.84 \pm 0.1^a$ | $0.39 \pm 0.05$ | $0.39 \pm 0.06$ | $0.44 \pm 0.03$ | $0.23 \pm 0.04^a$ |
| 0.4 | **$3.25 \pm 0.14^b$** | **$1.77 \pm 0.08^b$** | $0.27 \pm 0.03$ | $0.41 \pm 0.05$ | $0.43 \pm 0.04$ | $0.23 \pm 0.04^{ab}$ |
| 4 | **$3.33 \pm 0.17^b$** | $1.82 \pm 0.11^{ab}$ | $0.26 \pm 0.03$ | $0.41 \pm 0.06$ | $0.43 \pm 0.03$ | $0.23 \pm 0.04^{ab}$ |
| 40 | **$3.25 \pm 0.10^b$** | **$1.75 \pm 0.08^b$** | $0.26 \pm 0.02$ | $0.39 \pm 0.06$ | $0.43 \pm 0.04$ | $0.22 \pm 0.03^{ab}$ |
| 400 | **$3.27 \pm 0.46^b$** | **$1.80 \pm 0.18^b$** | $0.26 \pm 0.02$ | $0.38 \pm 0.06$ | $0.44 \pm 0.06$ | $0.23 \pm 0.03^{ab}$ |
| 4000 | **$3.27 \pm 0.2^b$** | **$1.78 \pm 0.1^b$** | $0.26 \pm 0.03$ | $0.39 \pm 0.06$ | $0.44 \pm 0.05$ | **$0.21 \pm 0.02^b$** |

### Behavior

Neither metformin (ANOVA DF=5, F=0.36, p=0.87; Fig. 3A) nor guanylurea (ANOVA DF=5, F=0.29, p=0.91; Fig 4A) exposed zebrafish had any alterations in the total distance traveled during a general swimming assay. There were no significant effects of metformin (ANOVA DF = 5, F =0.49, p = 0.78; Fig. 3B) or guanylurea (Kruskal-Wallis DF = 5, H =10.15, p = 0.071; Fig. 4B) exposure to thigmotaxis behavior. Metformin exposure did not affect total distance moved during dark (Kruskal-Wallis DF = 5, H =9.98, p = 0.076; Fig 3C) or light periods (ANOVA DF = 5, F =1.002, p = 0.42; Fig. 3D). Likewise, guanylurea exposure did not affect total distance moved during dark (ANOVA DF = 5, F =0.318, p = 0.902; Fig. 4C) or light periods (Kruskal-Wallis DF = 5, H =10.36, p = 0.066; Fig. 4D). There was no effect of metformin exposure on the acoustic response of zebrafish larvae at 120 hpf (Two-way ANOVA DF=5, F=0.63, p=0.67; Fig. 3E). Similarly, guanylurea had no impact on the response to acoustic stimuli (Two-way ANOVA DF=5, F=0.77, p=0.57; Fig. 4E).

**Fig 3:**
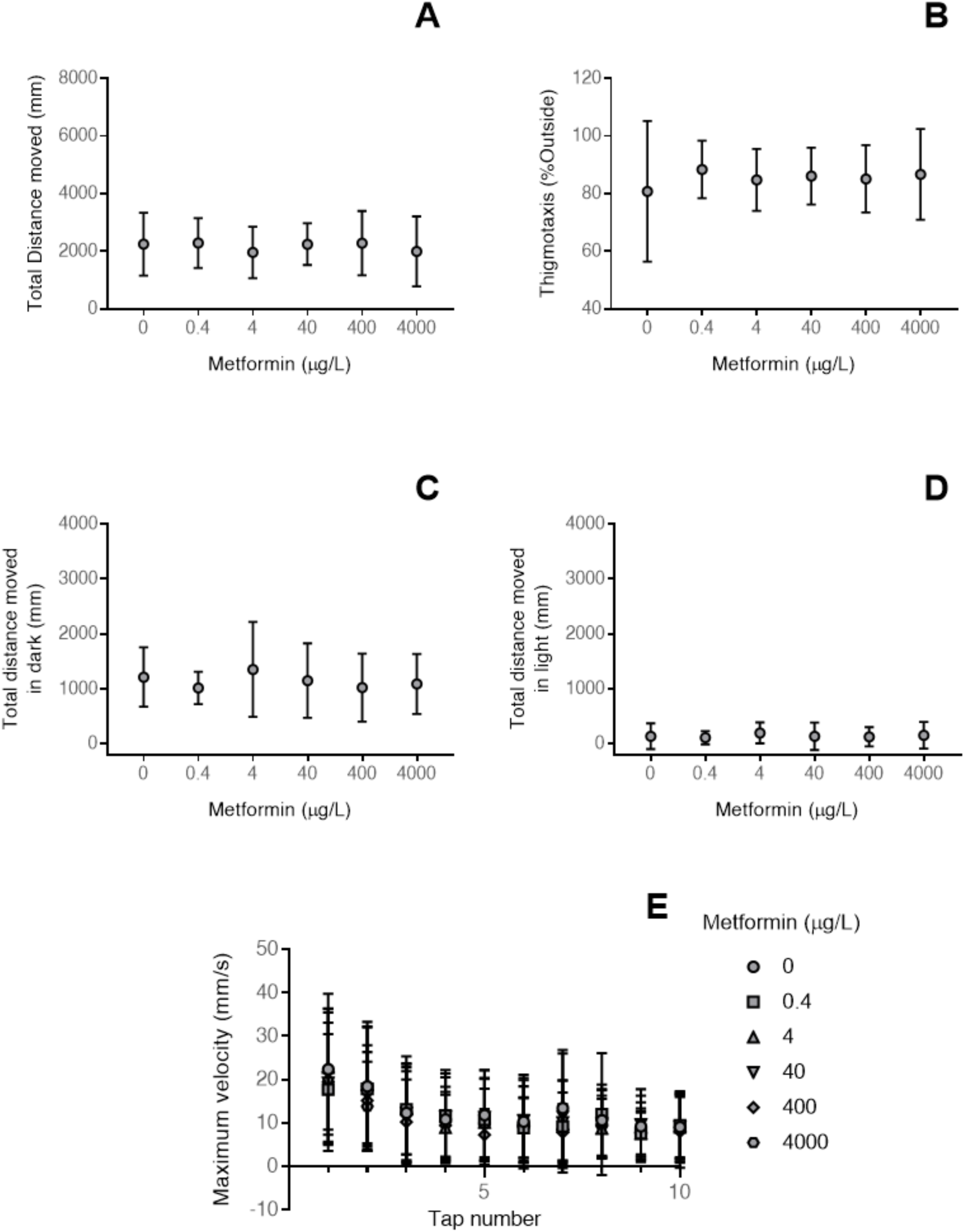
Developmental metformin exposure does not alter larval zebrafish behaviour. (A) The total distance travelled during a general swimming assay; (B) thigmotaxis; the total distance moved in the (C) dark and (D) light during a visual motor assessment; and (E) the maximum velocity following consecutive taps in larval zebrafish exposed to environmental and supra environmental levels of metformin throughout development. Values are displayed as mean ± standard deviation (n=16 for A and B; n=46-49 for C-E).

**Fig 4:**
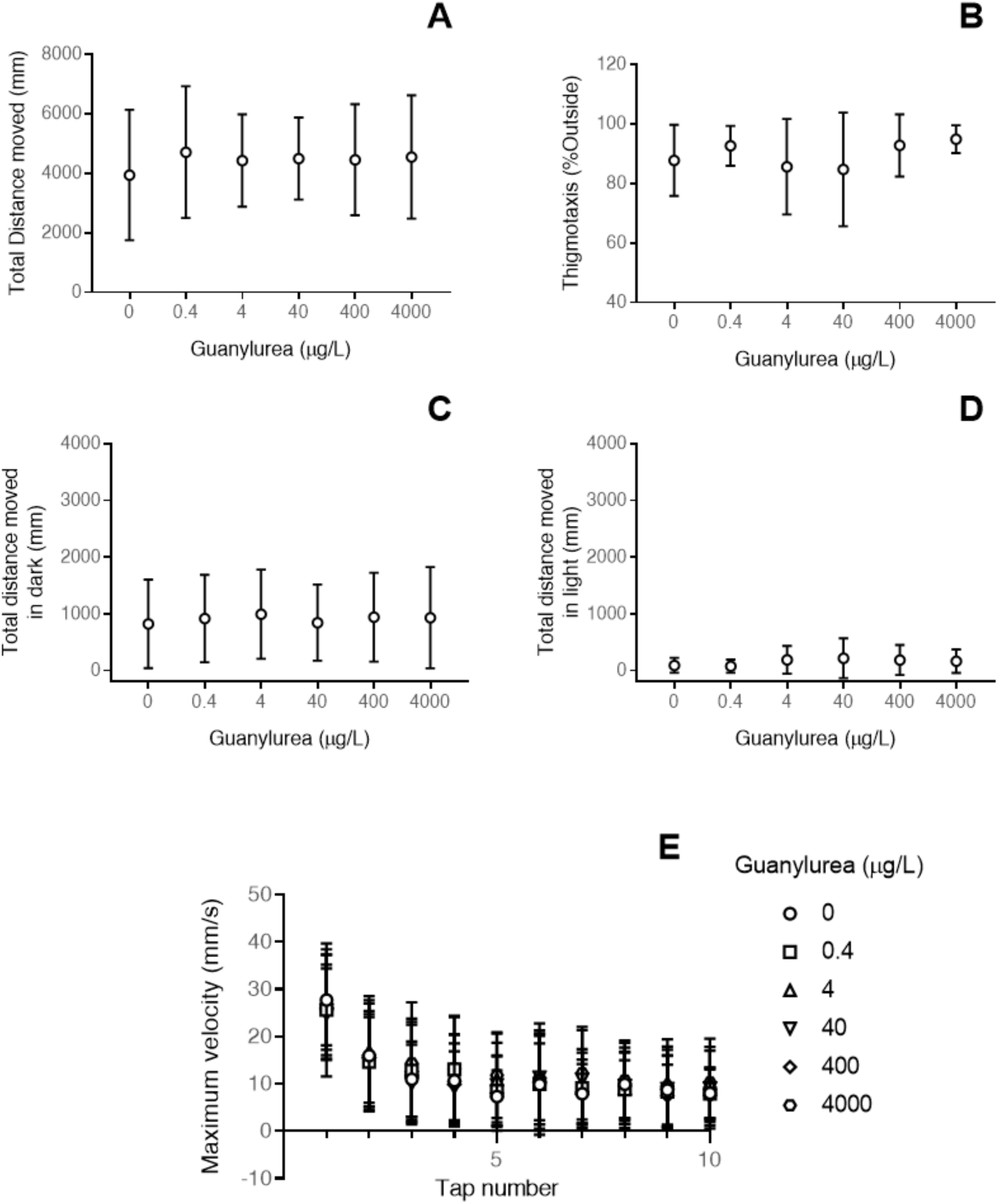
Developmental guanylurea exposure does not alter larval zebrafish behaviour. (A) The total distance travelled during a general swimming assay; (B) thigmotaxis; the total distance moved in the (C) dark and (D) light during a visual motor assessment; and (E) the maximum velocity following consecutive taps in larval zebrafish exposed to environmental and supra environmental levels of guanylurea throughout development. Values are displayed as mean ± standard deviation (n=16 for A and B; n=46-49 for C-E).

### Energetics

Oxygen consumption in zebrafish larvae exposed to metformin throughout development was not affected at 5 dpf (ANOVA DF=5, F=1.53, p=0.19; Fig 5A). Similarly, guanylurea exposure did not alter oxygen consumption of larval zebrafish (ANOVA DF=5, F=0.23, p=0.95; Fig 5B). There were no significant changes in glucose (ANOVA DF = 5, F =1.00, p = 0.43; Fig. 5C), glycogen (ANOVA DF = 5, F =1.68, p = 0.17; Fig. 5E) and ATP (Kruskal-Wallis DF = 5, H =6.69, p = 0.24; Fig. 5G) levels following exposure to metformin in larval zebrafish. Likewise, guanylurea exposure did not alter glucose (ANOVA DF = 5, F =1.82, p = 0.14; Fig. 5D), glycogen (ANOVA DF = 5, F =2.29, p = 0.072; Fig. 5F), ATP (ANOVA DF = 5, F =0.695, p = 0.63; Fig. 5H) levels in 5 dpf larval zebrafish. Glycogen levels appear to lower in our guanylurea exposure when compared to the metformin exposure. However, these exposures were conducted on separate embryos of zebrafish, at different times, and formal statistical comparisons were not carried out.

**Fig 5:**
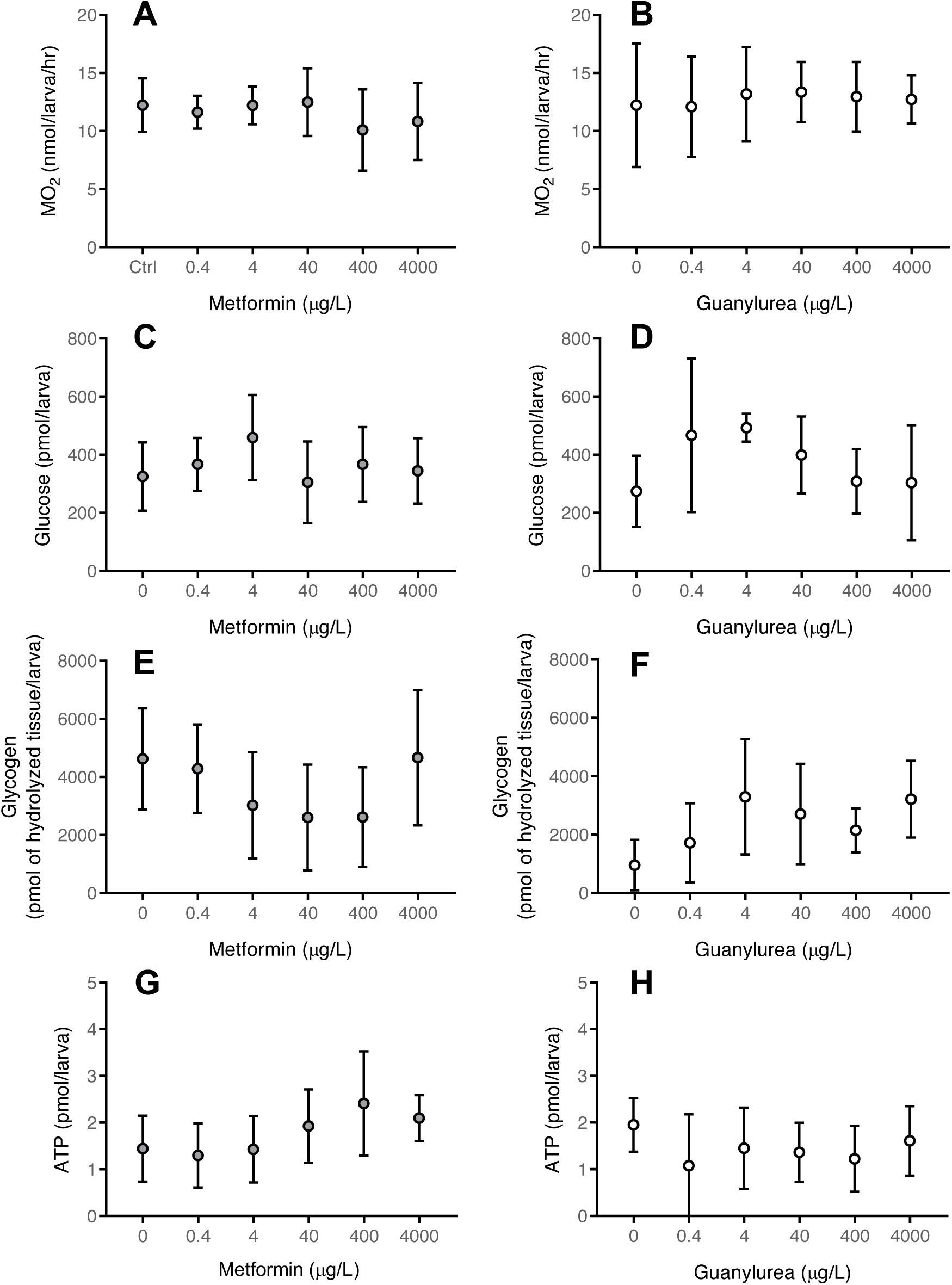
Exposure to metformin and guanylurea does not alter energetics at 120 hpf in zebrafish. The oxygen consumption, glucose, glycogen, and ATP levels of zebrafish larvae following exposure to metformin (A, C, E, and G, respectively) and guanylurea (B, D, F, and H, respectively). Values are displayed as mean ± standard deviation (n=12 for A and B; n=6 for C-H). Values are displayed as mean ± standard deviation (n=12 for A and B; n=6 for C-H).

### Heart rate

Metformin did not alter heart rate at 24 (Kruskal-Wallis DF=5, H=10.34, p=0.066; Fig. 6A) or 72 hpf (Kruskal-Wallis DF=5, H=9.069, p=0.106; Fig. 6E). However, at 48 hpf, metformin did decrease heart rate at 0.4 μg·L^−1^ (Kruskal-Wallis DF=5, H= 37.33, p<0.0001; Fig. 6C). At 24 hpf, guanylurea increased the heart rate of zebrafish at 4000 μg·L^−1^ (ANOVA DF=5, F=2.88, p=0.016; Fig. 6B). However, there were no effects in heart rate noted with guanylurea exposure at 48 (Kruskal-Wallis DF=5, H=7.1; p=0.21; Fig.6D) and at 72 hpf (Kruskal-Wallis DF=5, H=5.35, p=0.37; Fig. 6F).

**Fig 6.**
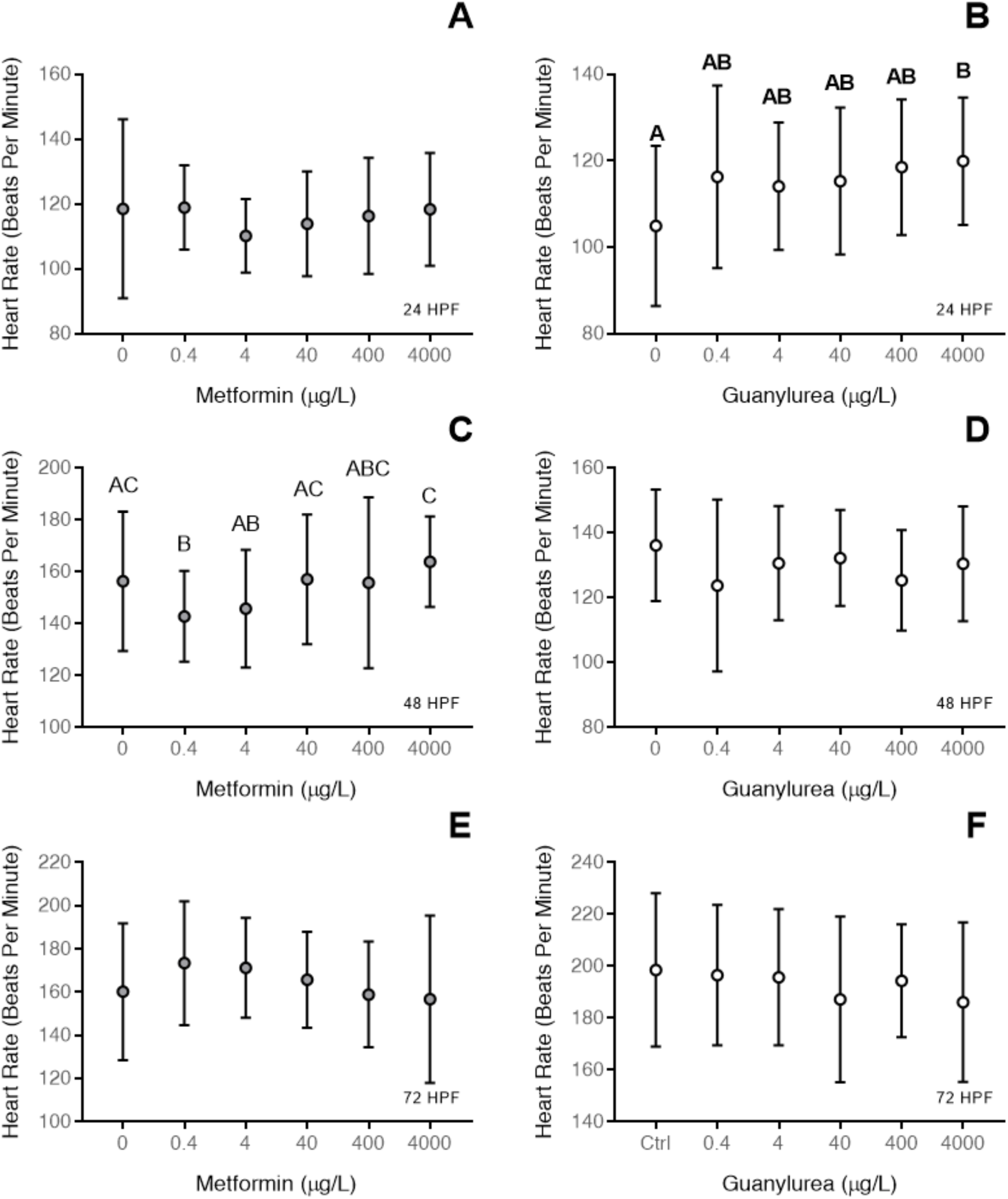
Exposure to metformin and guanylurea alters heart rate for short periods throughout early development. Heart rate at 24, 48, and 72 hpf after exposure to metformin (A, C, and E, respectively) and guanylurea (B, D, and F, respectively) throughout development. Values are displayed as mean ± standard deviation. Different letters denote significant differences between groups (n=45 for metformin; n=30 for guanylurea).

## DISCUSSION

The elevated occurrence of metformin (1-30 μg·L^−1^) in surface waters worldwide has raised concerns of the presence of this anti-hyperglycemic drug in the environment (Scheurer et al. 2012; Oosterhuis et al. 2013; Niemuth et al. 2015; Niemuth and Klaper 2015). While we assessed a gamut of performance and health markers in zebrafish, our results importantly indicate that exposure to concentrations just outside current environmental levels of metformin can increase the mortality rate of developing zebrafish. Studies in brown trout (Jacob et al., 2018), Japanese medaka (Ussery et al., 2018), and fathead minnow (Parrott et al. 2021; Parrott et al. 2022) have shown no increase in mortality following exposure to metformin in concentrations ranging up to 100 μg·L^−1^. However, our results support the recent studies in zebrafish larvae, where exposures to concentrations of metformin <100 μg·L^−1^ did increase mortality ((Elizalde-Velázquez et al. 2021; Phillips et al. 2021). Together, these increases in mortality may suggest that zebrafish are sensitive to metformin exposure compared to other fish species. However, significant sub-lethal indicators of effects, if any, were only seen at supra-environmental concentrations. We present some evidence that metformin can alter the normal growth of zebrafish, while increasing the incidence of malformations, but only at concentrations above the highest recorded in surface waters (≥40 μg·L^−1^). Together, these results suggest that metformin imposes a modest impact on developing zebrafish at environmental levels, leading to no overt changes in performance of fish entering feeding (120 hpf).

Early development is a tightly controlled and regulated process (Rougvie 2001). The developmental rate of an animal, then, would be an indicator of the level of controlled growth an animal exhibit. In the current study, the results suggest that metformin alters the normal growth patterning of zebrafish. Hatching, a process that typically begins in zebrafish at 48 hpf (Thompson et al., 2017), can be manipulated by temperature (Eme et al. 2015), but can be induced or delayed by contaminants (Thompson et al. 2017; Lee et al. 2019; Le Bihanic et al. 2020). Exposure to metformin at supra environmental levels (4000 μg·L^−1^) can delay the normal hatching window of zebrafish, suggesting a disruption to the development of these animals.

However, we have shown that *both* metformin and guanylurea have the capacity to disrupt the growth of zebrafish larvae, reducing the total length of the body and eye. The idea that these compounds have the capacity to impose abnormal growth on larvae is supported by the increased incidence of spinal curvature (>400 μg·L^−1^ metformin; 400 μg·L^−1^ guanylurea). However, the overt consequences of these changes in growth are unclear. For example, most effects are only seen at concentrations that are at, or beyond what has been recorded environmentally. Moreover, the effect size seen in total length may not be impactful for animal performance. While body length in fish has been correlated to swimming performance at the larval stage (Fisher et al. 2000) we did not observe any behavioural changes in any exposure treatment tested in this study. Together, our results suggest these alterations in growth may not be overly impactful for larvae, but whether continual exposure leads to persistent alterations in growth needs to be directly assessed.

In mammals, several studies have suggested that hepatic glucose production is directly affected following exposure to metformin (Rena et al. 2017), with the direct mechanisms unclear. For example, several studies have shown that metformin administration can directly alter mitochondrial respiration (Hawley et al. 2010) and inhibit mitochondrial glycerophosphate dehydrogenase (Thakur et al. 2018). This disruption to mitochondrial function leads to the activation of the cellular energy sensor AMP-activated protein kinase (AMPK), which acts to recover energy stores by altering cellular process to reduce ATP consumption and boost catabolic process that increase ATP production (Hardie 2011). In this study, we sought to test whether metformin exposure throughout development would lead to impairments in the energy status of zebrafish. However, our results suggest that metformin likely does not lead to disruptions in metabolism. This contention is supported by similar measures of oxygen consumption in fish exposed to environmental and supra environmental levels, a trend also seen in total ATP levels of larvae. The hallmarks of metformin action, decreased serum levels of glucose and reductions in gluconeogenesis (Hundal et al. 2000; Agius et al. 2020), are not reflected in our results. Given the size of larval zebrafish we were unable to measure plasma glucose levels, but overall, there appears to be no change in carbohydrate stores. This is surprising, given previous studies demonstrating clear reductions to COX (Barros et al., 2022), suggesting impairments towards ATP production. Perhaps, these results reflect metformin not posing an immediate threat to the metabolism of developing fish, and we must consider that these fish have not yet started exogenous feeding. Work has recently shown that a possible effect of metformin administration may be in reductions in glucose uptake at the level of the gut (Horakova et al. 2019), and repeating these measures in animals at more developed stages, such as following the onset of feeding, may be prudent to understand the possible effects of this anti-hyperglycemic drug in fish.

Behaviour may be sensitive to developmental and acute metformin exposures (Philips et al., 2021). In this study, we sought to apply a more extensive battery of testing to confirm disruptions in animal performance, which may represent substantial impacts to larvae in aquatic habitats (Fiksen et al. 2007). Therefore, we applied a general swimming assay, a light-dark visual motor paradigm, an acoustic response test, and a measurement of larval thigmotaxis, to understand the implications of metformin exposure to animal function. The light-dark paradigm is a well-established hyperactivity model in zebrafish, with the transition to dark following a prolonged exposure to light in 120 hpf zebrafish inducing a period of increased activity (Thompson et al., 2017). To our knowledge, this is the first assessment of thigmotaxis following exposure to metformin and guanylurea in fish. Changes in thigmotaxis, or the anxiety-like behavior in fish, may have large implications in ecological settings, influencing the capacity of fish to capture prey, explore novel environments, or as has been shown in *Gammarus fossarum,* can improve predator evasion (Perrot-Minnot et al. 2017). General swimming assays can offer insight into the baseline activity of an animal in an undisturbed environment (Fisher and Bellwood 2003). The acoustic startle response has been suggested to be a basic form of learning, exemplified by the cessation of the animal’s response to repeated signal (Beppi et al. 2021), and has been proposed to be representative of avoidance behaviour (Faria et al. 2019). The application of these assessments demonstrates that metformin exposure does not impart any changes to animal performance, light perception, or rudimentary learning. This is in accordance with prior studies that have shown no impacts to the general swimming capacity of juvenile brown trout (Jacob et al., 2018) and in zebrafish larvae exposed to a similar light-dark paradigm (Godoy et al., 2018). Another study using zebrafish demonstrated a larger hypoactive response following metformin exposure from 4 hours post-fertilization until 5 dpf (Philips et al., 2021). However, the results from the aforementioned study were taken from AB strain zebrafish (Philips et al., 2021), and perhaps the negative results of the current study are a product of a strain-specific effect. A tempting speculation would be that metformin also does not impact neurodevelopment in fish, as behavioural alterations have been proposed to be an indicator of neurotoxicity (Tierney 2011), with recent studies tying behavioural disruptions to changes in neurodevelopment following contaminant exposure (Kinch et al. 2015; Thompson et al. 2017; Thompson and Vijayan 2020b). However, this remains to be explicitly tested following early-life metformin exposure.

Considering the mode of action of metformin, we sought to investigate cardiometabolic parameters in these fish. After uptake, metformin accumulates in the hepatic inner mitochondrial membrane where it inhibits NADH-ubiquinone reductase (mitochondrial complex I), preventing mitochondrial ATP production, increasing ADP: ATP and AMP: ATP ratios (Luengo et al. 2014). Through complex 1 inhibition, NADH oxidation is decreased, inevitably decreasing oxygen consumption in mammalian systems (El-Mir et al. 2000). Metformin has been shown to elevate genes such as smdt1a, atp2a2a, tnnilc, and hsd11b2 (10 000 μg·L^−1^), which have been implicated in cardiovascular development and function (Philips et al., 2021). Indeed, our results suggest that metformin exposure can influence the early function of the heart (as inferred via heart rate). We see changes in heart rate at 48 hpf; however, the differences noted are not dose dependent, and the effect itself is mildly bradycardic. Little evidence exists directly tying changes in heart function to metformin exposure (Manzella et al. 2004), despite the advantages to endothelial function, protection from oxidative stress, and hypertrophy induced via angiotensin II (reviewed by Nesti and Natali, 2017). This may suggest that our results are not a product of the direct interaction of metformin with the heart, but a development effect. A recent study has suggested that antidepressants can lead to precocious development of the heart, reflected in a more advanced heart rate (Thompson et al. 2022), and perhaps a diminished heart rate is a representation of a delayed development. The changes in heart rate in this study are not mirrored in our measurements of oxygen consumption, possibly because oxygen consumption was measured much later (120 hpf) in development, but also because aerobic metabolism in zebrafish is independent of the cardiac system in early-life stages (Pelster and Burggren 1996). Therefore, longer-term studies assessing cardiac function may be necessary, particularly exploring developmental windows where zebrafish transition away from transcutaneous respiration (Burggren 2004).

Guanylurea, the major bacterial transformation product of metformin (Caldwell et al. 2019), can be found at higher levels in effluent than its parent compound (Scheurer et al. 2012). A major aim of this study was to assess the sublethal impact of this metabolite relative to metformin. Similar to metformin, guanylurea increased mortality in larval fish and elevated spinal curvature. While spinal curvature appears to occur at supra-environmental levels, mortality was increased in concentrations that are just outside the maximum range of values reported currently (28 µg·L^−1^; Scheurer et al., 2012). Guanylurea, however, may appear to be more impactful in disturbing the normal growth of larvae, as environmentally relevant levels lowered the total size of larvae. A similar result was seen in Ussery et al. (2019), where guanylurea imposed limitations in growth at the ng·L^−1^ level compared to impacts of metformin seen in µg·L^−1^ exposures. A recent study has positioned guanylurea as a more toxic compound to zooplankton than metformin (Gao et al. 2023), but the results of this study suggest that guanylurea and metformin are more or less equivalent in their impact to larval development.

In conclusion, the results of the current study reveal a modest impact of metformin and guanylurea to zebrafish larvae exposed to these compounds from 4 hpf. While we did indeed observe increases in mortality, this occurred at concentrations currently beyond the extreme reported environmentally (Scheurer et al. 2012; Oosterhuis et al. 2013; Niemuth et al. 2015) and the effect size was small (∼5-10% increase). Further, while our results demonstrate that metformin has the capacity to alter the development, growth, and heart rate of zebrafish, the concentrations needed to produce these disruptions are similarly supra-environmental.

Metformin can influence feeding responses in mammals (Aubert et al. 2011), and improve glucose removal in the plasma of rainbow trout fed on a high carbohydrate diet (Panserat et al. 2009). Indeed, an exposure with Japanese medaka demonstrated a clear growth retardation following longer-term exposure (28 day old) that was not evident at earlier life stages (Ussery et al., 2019), which may be suggestive of impairments becoming observable following the transition to external energy sources. As our study design did not encompass the early feeding window for this species, future investigations targeting the transition to exogenous feeding may be necessary to understand limitations to feeding, and absorption, following exposures to metformin and guanylurea.

### CrediT authorship contribution statement

Conceptualization: JYW, SW, WAT, OB; Investigation: SW, WAT, NM, ME, JQ, OB; Formal Analysis: WAT, SW; Writing-original draft: SW, WAT; Writing-review and editing: SW, WAT, OB, ME, JQ, NM, JYW; Supervision: JYW, SW, WAT, OB; Funding: JYW

## Acknowledgements

We are grateful for the help of Lisa Laframboise and Dr. Derek Alsop for their efforts in maintaining the lab populations of zebrafish. This study was supported by the Natural Sciences and Engineering Research Council grant awarded to JYW and a NSERC Postdoctoral fellowship awarded to OB.

